# Recapitulating tumor microenvironment using preclinical 3D tissueoids model for accelerating cancer research and drug screening

**DOI:** 10.1101/2020.12.21.423825

**Authors:** Ambica Baru, Saumyabrata Mazumdar, Prabuddha Kundu, Swati Sharma, Biswa Pratim Das Purakayastha, Sameena Khan, Reeshu Gupta, Nupur Mehrotra Arora

**Affiliations:** Mammalian Cell Culture Lab, Premas Biotech Pvt Ltd., Sector IV, IMT, Manesar, Gurgaon-122050

**Keywords:** Tissueoid, 3D/4D culture, Cancer, Drug Discovery

## Abstract

The formation of three-dimensional spheroid tumor model using the scaffold-based platforms has been demonstrated over many years now. 3D tumor models are generated mainly in non-scalable culture systems, using synthetic and biological scaffolds. Many of these models fail to reflect the complex tumor microenvironment and do not allow long-term monitoring of tumor progression. This has resulted in inconsistent data in drug testing assays during preclinical and clinical studies. To overcome these limitations, we have developed 3D tissueoids model by using novel AXTEX-4D™ platform. Cancer 3D tissueoids demonstrated the basic features of 3D cell culture with rapid attachment, proliferation, and longevity with contiguous cytoskeleton and hypoxic core. This study also demonstrated greater drug resistance in 3D-MCF-7 tissueoids in comparison to 2D monolayer cell culture and the collagen-based 3D system. In conclusion, 3D-tissueoids are more responsive than 2D-cultured cells in simulating important tumor characteristics, anti-apoptotic features, and their resulting drug resistance.

## Introduction

Cancer is a global health issue that continues to be a challenge to treat and demand action^1^. The development of effective anticancer drugs significantly requires and depends on reliable in vitro high-tech screening systems^2^. The absence of reliable and effective in vitro screening models that could mimic key aspects of the tumor microenvironment, such as drug resistance and phenotypic changes to cells, impedes the reliable translation of in vitro findings into in vivo clinical models. The poor correlation between preclinical *in vitro* and *in vivo* data with clinical trials has an adverse impact on drug development costs, with the trial cost vary from $2 million to $347 million^3^. Only ∼7% of anticancer drugs gain clinical approval, much lower than drugs for other diseases^4^. Therefore, the action is required to develop effective and reliable in vitro models that reflect the *in vivo* tumor microenvironment and *in vivo* efficacy more accurately. Two-dimensional (2D) monolayer cell culture has contributed immensely to understanding basic cancer research, disease modelling, drug discovery, and toxicity studies. However, 2D cell culture-based assays are mostly non-indicatory, non-predictive, and non-representative of the real tissues^5,6^. To this end, 3-D tumor spheroids are an attractive alternative to 2-D cell culture as they can recapitulate many aspects of the tumor microenvironment, including paracrine effects, cell-cell interactions, and extracellular matrix deposition^7,8^. Furthermore, many environmental factors inducing metabolic and oxidative stress in tumor cells such as hypoxia and the formation of a necrotic core can also be recapitulated by 3D systems^9^. Additionally, 3-D spheroids also reduce the time and costs associated with translating laboratory findings into animal models and can accelerate the process of anticancer drug development^2^.

The establishment of 3D cell culture methods has been based on either scaffold-based, scaffold-free gels, bioreactors/or microchips^10^. Various scaffold-based platforms have been described in the literature, which can be biological (hydrogels such as collagen, gelatin, alginate, or chitosan), synthetically engineered (Polyethyleneglycol (PEG), polylactic acid (PA), polyglycolic acid (PGA)) or solid scaffolds to emulate key properties of ECM. However, these systems are not entirely efficient in being tedious to produce, time-consuming, unstable over a long period, and may pose sample retrieval challenges^10-13^. Moreover, a synthetic scaffold-based 3D system allows cells to spread in a limited fashion and does not resemble the 3D cell culture characteristics^14^ and may have flattened morphology. Similarly, scaffold-free systems, often referred to as spheroids, can be produced in various ways, such as the hanging drop technique. However, the routine use of such models for drug development has been hampered by a lack of standardized procedures to produce uniform spheroids and high-throughput screening^15^. Therefore, the lack of a robust 3D cell culture model led to the continued improvement in the 3D culture system and technology.

The study addresses the existing challenges in the prior art to achieve the functional outcome by utilizing a three-dimensional tissueoids model that closely recapitulate the tumor microenvironment. The model has been developed on AXTEX-4D™ platform that removes the need for additional biomolecules(such as hydrogels) and instead relies on the cells themselves to create their own gradients and microenvironmental factors^16,17^. The fabric of AXTEX-4D™ includes non-woven fabric, porous, an inert polymer that allows the cells to attach, grow into its spaces and, help in acquiring 3D like features^16,17^.

In this study, cancer tissueoids demonstrated the basic features of 3D cell culture, such as universality, rapid attachment, longevity, viability, hypoxia, and cytoskeleton arrangement, which are required for the 3D growth of cell lines and biopsies. We have further compared the efficacy of drug sensitivity with other 3D cell culture models and 2D-monolayer culture. Overall, the current study characterizes and highlights the utility of the preclinical 3D tissueoids model, which may be useful for analyzing features of growth and drug sensitivity of cancer cells.

## Results

### Universality: Formation of 3D tissueoids from various mammalian cell lines

The universality of 3D culture systems is particularly attractive as cells of different origins could be grown to enhance cancer research and drug development significantly. Therefore, we decided to define the universality of AXTEX-4D™ platformby growing cancer and transformed cells of different origins. This study used seven mammalian cell lines (cancer: MCF-7, PC3, Hep-G2, A-375; transformed: HEK-293, NIH-3T3, CHO-K1). Approximately one day after seeding on AXTEX-4D™ platform, tissueoids were formed by all seven cells. Previous findings have shown that 40 tumor cell lines of diverse origin, when cultured in 3D spheroid conditions, form two distinct groups according to the architecture of spheroid shapes: i) tight spheroids and ii) loose spheroids^18,19^. To this end, 3D tissueoids assembly and their growth on AXTEX-4D™ platform was determined by their shape and compact nature using scanning electron microscopy (SEM). Four (HEK-293, NIH-3T3, MCF-7, and CHO-K1) of the seven cell lines started to form compact tissueoids (CTs) (Fig 1A). For PC3, Hep-G2, and A-375, the cells grow in the AXTEM-4D™ platform, and although tissueoids formed, they were loose than those produced by the other four cell lines(Fig. 1B). These data support that tissueoids developed on AXTEX-4D™ showed 3D tissue-like organization, cell-cell connection, and interaction with the scaffold (Fig. 1A-1B). Interestingly, patterns like grooves and folds which mimic the real tissue morphology were observed in all 3D tissueoids. These data collectively indicate that cell growth in the 3D tissueoids model allows the cells to retain cellular organization, resembling the *in vivo* condition more closely.

**Figure 1.**
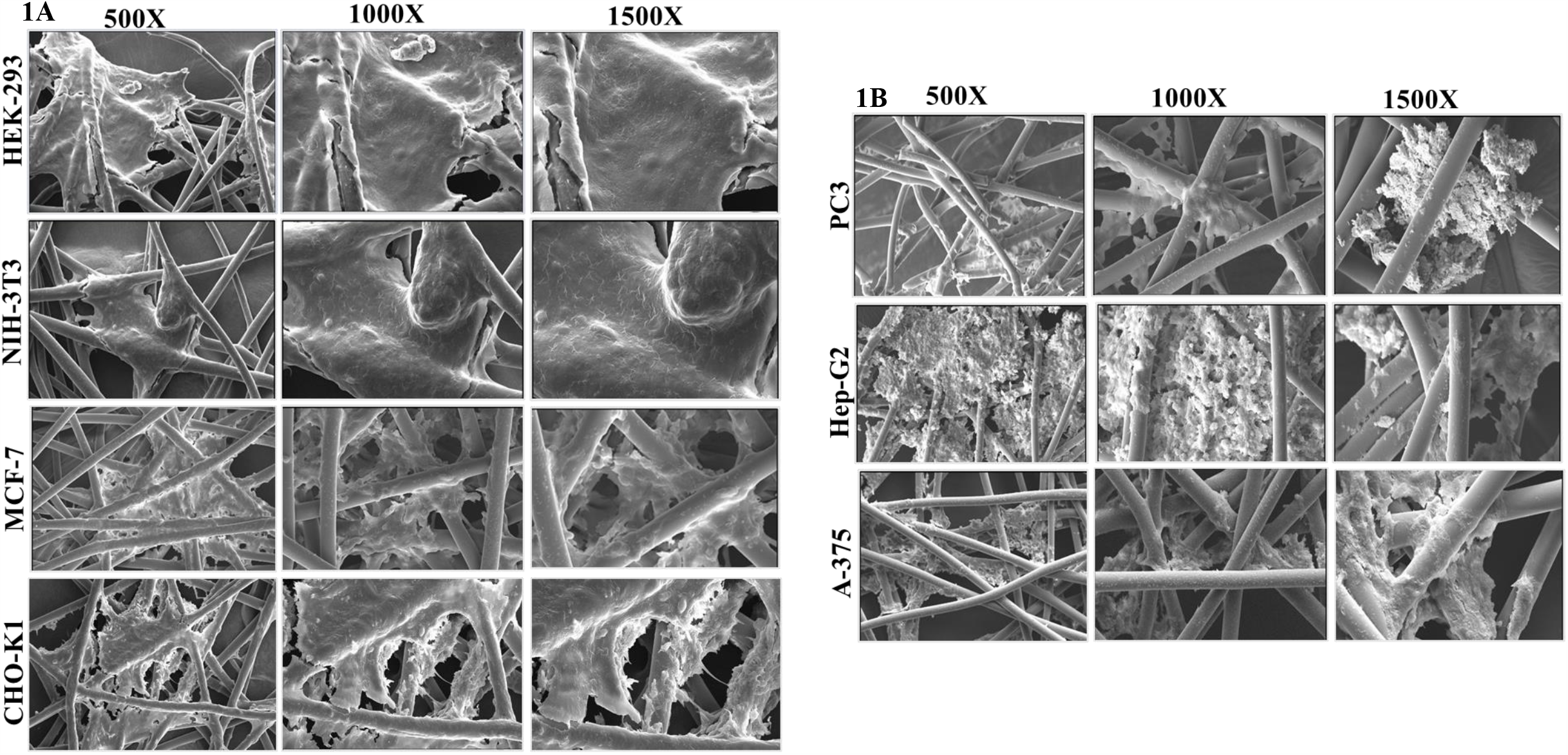
Formation of compact and loose 3D tissueoids on AXTEX-4D™ platform. Scanning electron microscope (SEM) images of A) compact and B) loose 3D tissueoids of indicated cell lines on the AXTEX-4D™ platform at 500X, 1000X and 1500X magnification

### Tissueoids formation with different number of plated cells

The ability to generate spheroids from small cell numbers is particularly relevant when dealing with rare patient-derived cells or cells with high mortality rates^20,21^. To observe the effect of cell density on tissueoids formation, different cell numbers of MCF-7 tissueoids were added to each well in a 96 well plate. To this end, SEM analysis showed that a plating density of even 25 cells/well resulted in consistent spheroid size and shape, with sizes suitable for image acquisition and analysis (Fig 2A). At this density, the average tissueoid maximum diameter was consistent with a value of 55 ± 5μm (n = 96), yielding a coefficient of variation of 9.0%. To achieve reproducible results in drug discovery, size uniformity is demanding since it affects cell behavior, function, drug penetration and efficacy, nutrient, and oxygen transport. Next, the dependence of the tissueoids size on the number of plated cells using the MCF-7 cell line was studied using phase-contrast microscopy. Obviously, the larger the original cell sheet, the bigger the 3D tissueoids obtained from the MCF-7 cell line. Our results support the obvious previous findings where it has been shown that increasing the density of cell seeding resulted in a linear increase in spheroid diameter^22,23^ (Fig 2B-2C). These results suggest the adaptability of the AXTEX-4D™ platform in forming 3D tissueoids when the sample availability is minimal. We have also tested the scaffold reproducibility in forming the tissueoids with different plate formats from 12, 48, and 96 well plate and found consistency with all the plate format suggesting the versatility of the AXTEX-4D™ platform to form tissueoids for desirable experimental needs (Fig 2D).

**Figure 2.**
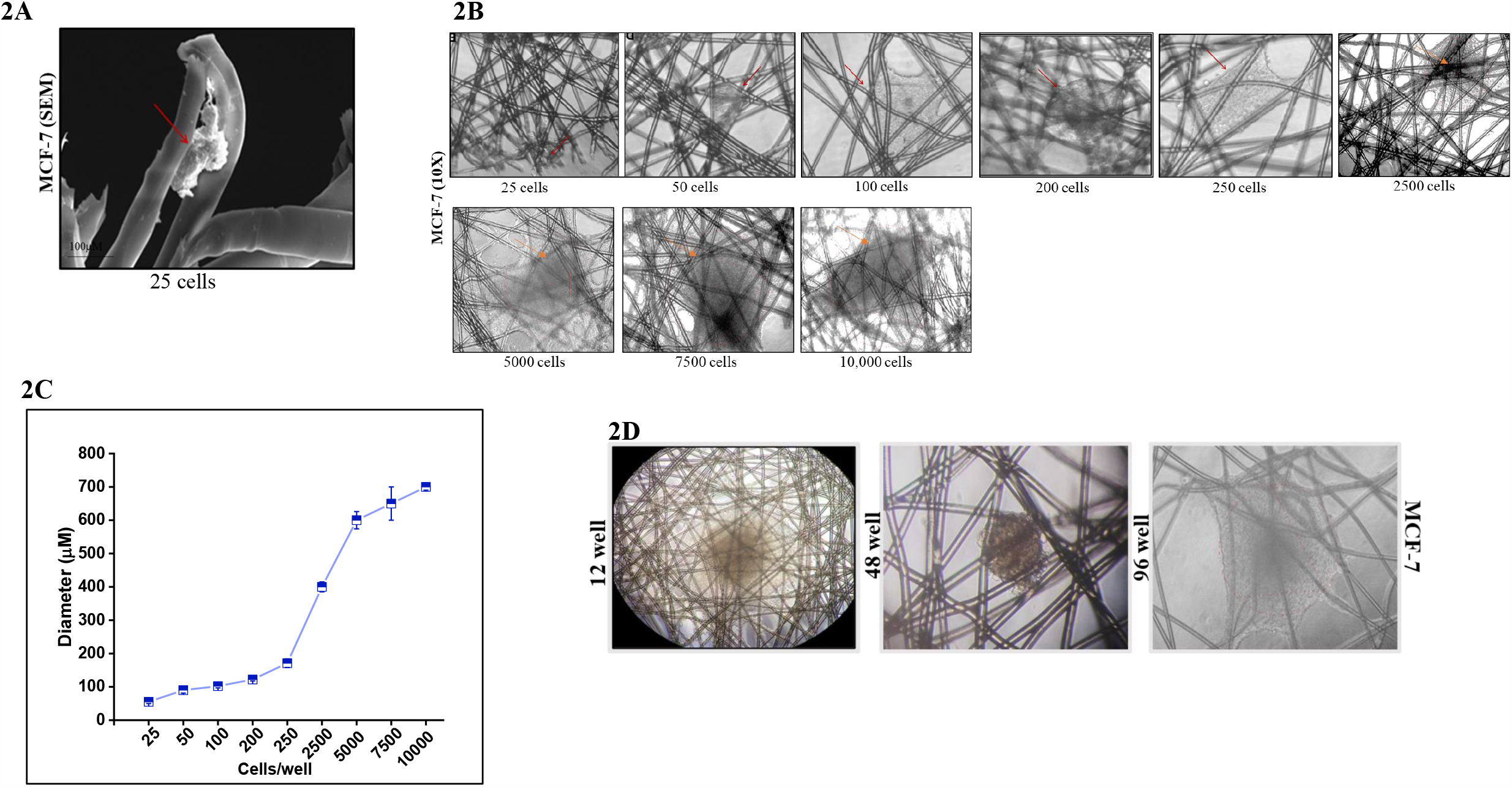
Diameters of 3D tissueoids with various cell densities. A) SEM image of MCF-7 3D tissueoids formed with 25 cells/well. B) Phase-contrast microscope images showing the diameter of MCF-7 3D tissueoids formed with indicated cell densities C) Graphical representation of diameters formed with various cell densities. Values are means ± S.E.M. from n=3 D) Phase-contrast microscope images showing the formation of MCF-7 3D tissueoids in indicated cell formats

### Rapid attachment of 3D Tissueoid

Rapid attachment of 3D spheroids may enhance high throughput drug testing and screening^24^. To this end, eight different cell types of different origins were tested for tissueoids formation in AXTEX-4D™ platform. Data of three representative cell lines (PC3, HEK-293, and NIH-3T3) are shown in this study. All the three tumor cells tested gave rise to compact, rigid, and spherically shaped tissueoids by 48 h after cell plating. For PC3 cells, it took nearly 48 h for these clusters to form tissueoids (*i*.*e*., clusters which are not dislodged by pipetting). However, tissueoids formed in less than 24 hours for the other two transformed cell lines (Figure 3A). These results suggest tissueoids generation within 24-48 hours by all the cell lines, which may enhance the practical applicability of the 3D tissueoids model.

**Figure 3.**
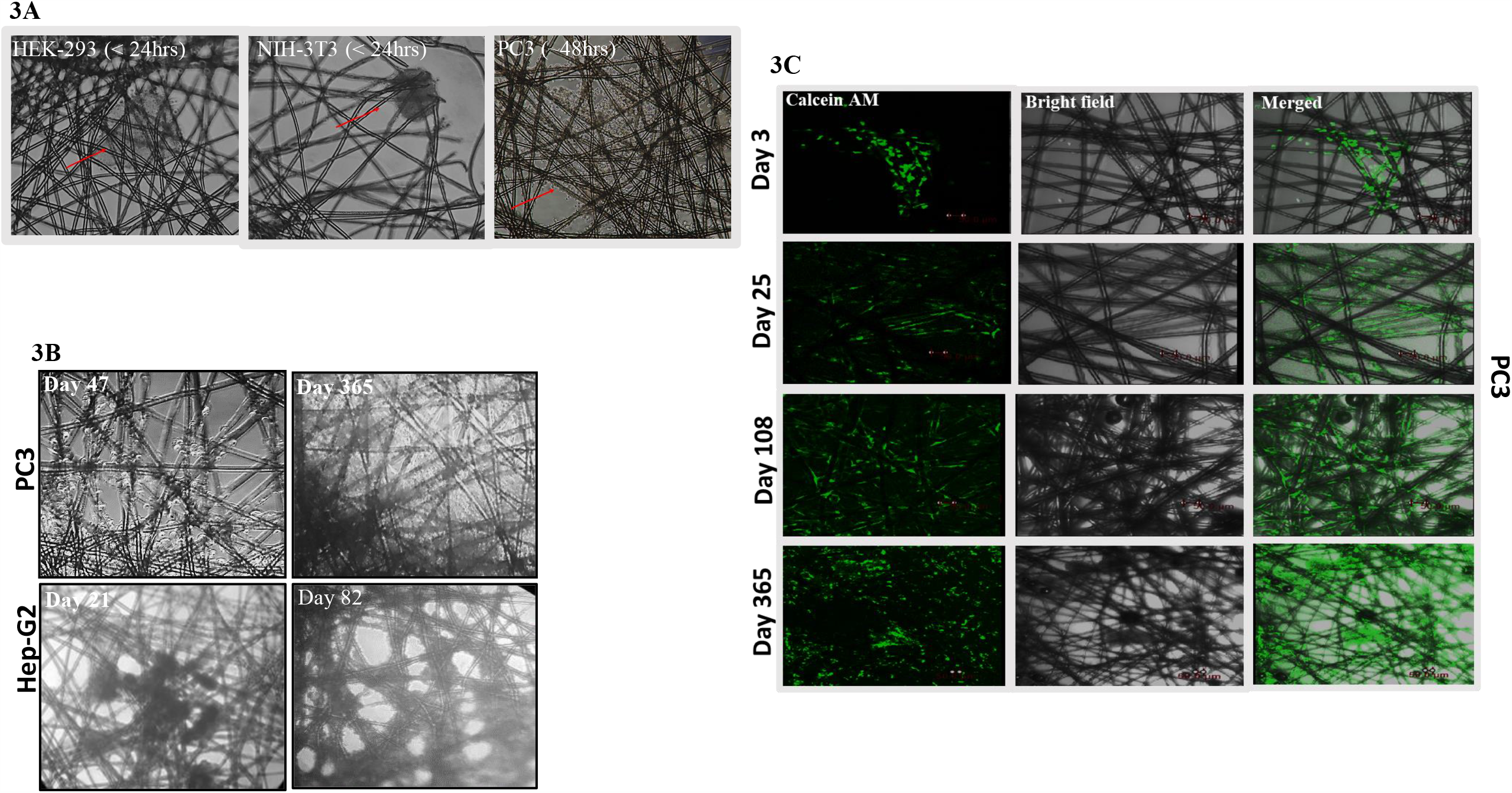
Attachment, longevity and viability of 3D tissueoids. 3D tissueoids of indicated cell lines were grown on AXTEX-4D™ platform and phase contrast microscope images at 10X magnification showing A) attachment and B) longevity were taken C) Confocal microscope images of PC3 3D tissueoids stained with Calcein AM showing viable cells at indicated time points

### Longevity and viability of 3D tissueoids

Longevity of the spheroid, a crucial attribute, is required for increasing flexibility and robustness of the 3D system to design the experiments^25,26^. To this end, the longevity and viability of the tissueoids were observed in two cancer cell lines: Hep-G2 and PC3. To show the length in time during which tissueoids remained in culture, the longevity of tissueoids derived from HepG2 and PC3 cells was observed. Tissueoids were generated in 12 well formats with media change after every third day. Different fields on different days were captured. This study demonstrates that tissueoids of HepG2 and PC3 were able to survive till 82 and 364 days with an increased number of cells and density, hence allowing the fourth dimension to the tissueoids. The fourth dimension is provided by the AXTEX-4D™ platform to generate the 3D tissueoids for an extended period of time^16^. We speculate that the smaller life span of Hep-G2 cells may be due to their high metabolically active nature and due to which these cells grow rapidly and, after a certain period, come out of the matrix ^27^(Fig. 3B). On the other hand, prostatic cell metabolism does not conform to the “classical” Warburg metabolic phenotype exhibited by other solid tumors^28^. These results suggest the applicability of the platform for time-dependent studies, single and multi-drug studies. Next, the viability of the tissueoids (PC3) at different time points (5, 25, 108, and 365 days) was analyzed by Calcein AM staining, which is used to determine cells’ viability. Cells showing green fluorescence were considered as viable. We have observed an increase in fluorescence over the period suggesting the long-term viability of tissueoids (Fig 3C).

### Cytoskeleton arrangement and hypoxic core in 3D Tissueoids

The dimensionality of a cell’s environment greatly influenced the structure and distribution of the cytoskeleton^29^. Therefore, in this study, we aimed to determine whether the 3D tissueoids model demonstrates this arrangement and compare it with 2D monolayer culture. Confocal analysis showed significant changes in the architecture of 3D MCF-7 and PC3 tissueoids, where contiguity of the cells was visible. In contrast, the cells in 2D were elongated and scattered with defined edges and margins (Fig. 4A & 4B). Furthermore, multiple hypoxia-driven cell behaviors and phenotypes reflect key characteristics of *in vivo* growing tumors^30^. We observed that HIF-1 α fluorescence was significantly higher in the core than the periphery of the 3D tissueoids for MCF7 cells when compared with 2D monolayer culture. These results suggest that 3D tissueoids have characteristics more similar to *in vivo* tumors than 2D monolayer culture.

**Figure 4.**
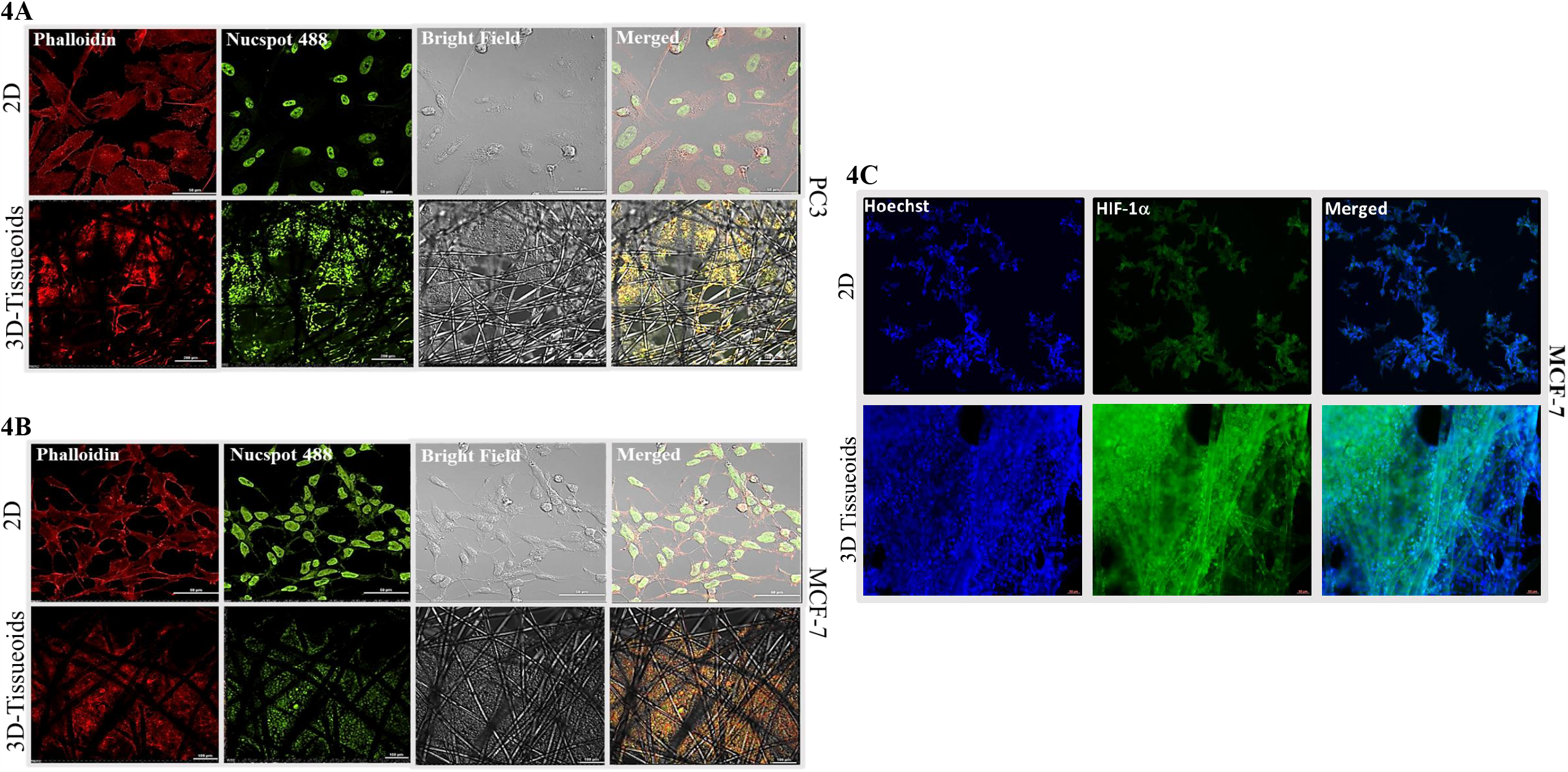
Cytoskeleton arrangement of 2D and 3D tissueoids. Confocal microscope images depicting cytoskeletal arrangement in 2D monolayer and 3D tissueoids using (A) PC3 and (B) MCF-7. Staining was performed with anti-Phalloidin antibody (Red) and Nuncspot Live 488 (green-nucleus stain) C) Immunofluorescence of MCF-7 cells depicting HIF-1α expression (green) in 2D and 3D tissueoids

### Effect of doxorubicin on viability and apoptosis

To explore the mechanism of differential sensitivity to doxorubicin in the 2D- and 3D-cultures, the viability of the MCF-7 cell line was compared in two 3D models: tissueoids and collagen. Specifically, 48-hour treatment with five doses (0.5, 1, 2.5, 5, and 10μM) of doxorubicin was carried out in 96 well plate. Results of prestoblue staining indicated that the MCF-7 3D tissueoidsthat developed on AXTEX-4D™ platform tended to show relatively high drug resistance at every dose as compared to the 2D and collagen model (Fig. 5A). We substantiate this data using Annexin/7AAD by FACS analysis (Fig. 5B). The data showed that viability of 3D cells decreases in a dose-dependent manner with ∼60-70% viability at high dose (10µm) in 3D tissueoids and collagen 3D while only ∼17% of cells were viable in a 2D monolayer at this dose (2D: 0.5µm-22±2%; 1µm-23±1.4%; 2.5µm-24±3.4%; 5µm-28±5%; 10µm-17±2.4%). Further, necrosis at a higher dose (10µm) of doxorubicin could be seen in both the 3D cultures (3D tissueoids: 15±0.3%; collagen: 21±5%), however, marginal apoptosis was observed in 3D tissueoids and collagen models (Fig 5C). These results suggest that 3D tissueoids could be an optimal drug-testing model, especially for analysis of long-term drug regimen and repeated doses.

**Figure 5.**
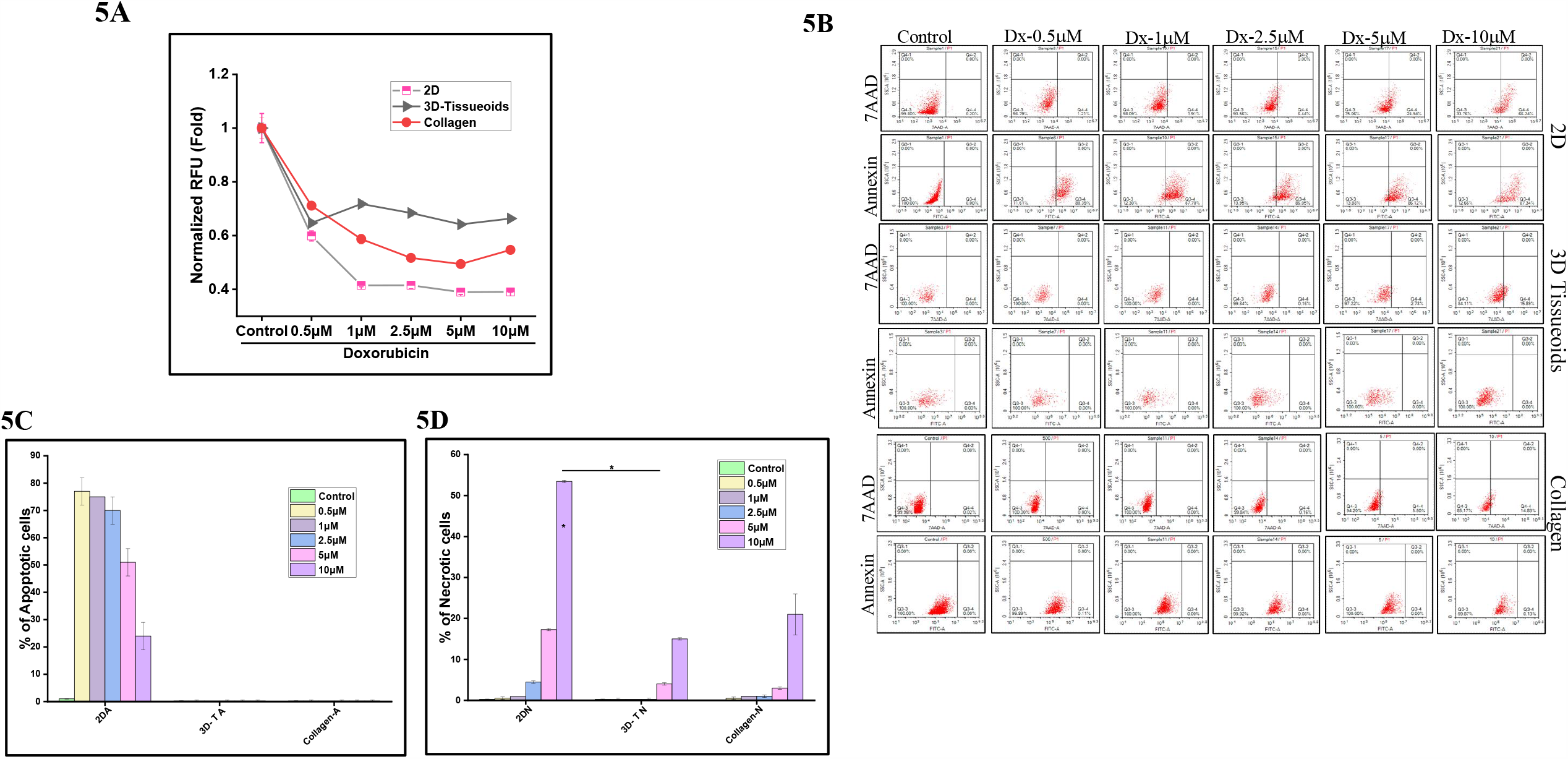
Effect of doxorubicin drug sensitivity in conventional 2D and indicated 3D cell culture models. MCF-7 3D tissueoids were treated with indicated concentrations of doxorubicin for 48hrs and (A) cell viability using PrestoBlue (B) apoptosis by using annexin V and necrosis by using Annexin-7AAD staining in flow cytometry was measured. Graphical representation of C) apoptosis and D) necrosis induced by various doses of doxorubicin. T=tissueoids; A=Apoptotic, N=Necrotic. Values are means ± S.E.M.

**Figure 6:**
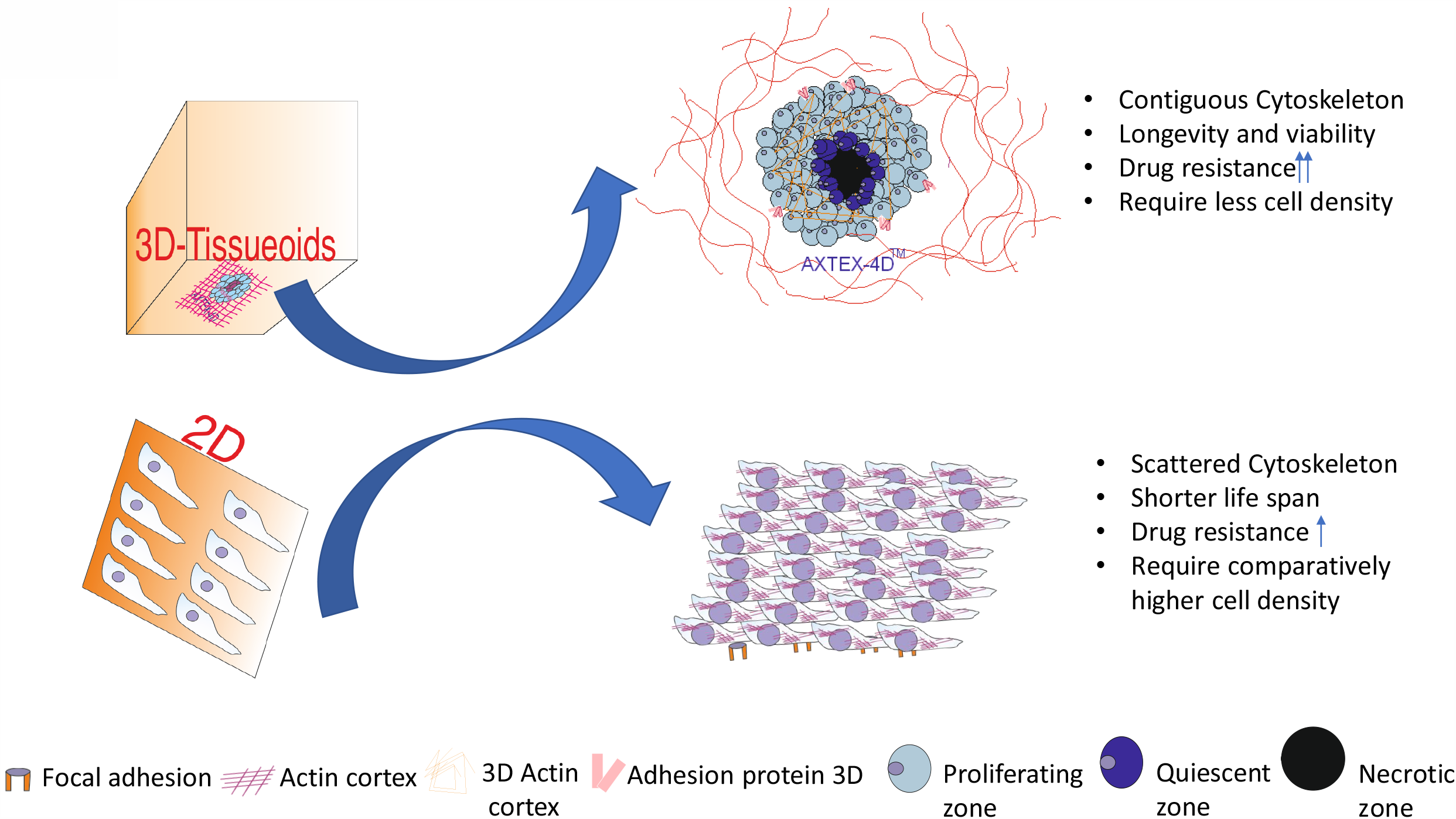
Schematic representation showing morphological differences in 2D monolayer culture and 3D tissueoids

## Discussion

In the last decade, multiple attempts have been made towards the improvement of preclinical models to accelerate cancer research. Despite these efforts, most of the available models fail to recapitulate several important attributes of the tumor microenvironment and do not reflect the complexity of human tumors^31,32,2^. However, against the optimism of these exciting possibilities, critical challenges for 3D cell cultures include assay validation, poor reproducibility between batches of scaffolds, and correlation to historical 2D culture results. Besides, most in vitro experiments can only be carried out for short culture periods, which impedes examination of long-term effects of the microenvironment or drug treatments on disease progression^33^.

In this work, we aimed to develop a 3D tissueoids model that is simple, ready to use without the need for sophisticated or complex infrastructure. The 3D tissueoids model was developed by using AXTEX-4D™ platform that allows long-term culture with continuous monitoring^16,17^. This platform uses a non-woven fabric scaffold to create 3D tissue-like structures with a function of time^16,17^. A fabric of the platform permits the cells to attach and allows sufficient permeability of gases & nutrients during culture. Cells grow across the scaffold over time which provides the cells a 3D tissueoids structure. Tumour-stroma crosstalk in 3D tissueoids was achieved by keeping cells in proximity and by accumulation of extracellular matrix components^16^. 3D tissueoids possessed cellular morphology with evident actin cytoskeleton arrangement along with hypoxic core and showed longer lifespans compared to their counterparts cultured on the standard 2D monolayer. The longer life span of tissueoids (∼365 days) makes them a suitable model for long-term cell cultures and allows for time-dependent studies, drug combinations, and multi-dosing studies with the potential for developing novel therapeutics. In contrast to this, other time-intensive 3D models based on collagen, matrigel, etc., have been reported to have a short life span due to the unavailability of the restricted flow of nutrients and gases^13,34^.

Furthermore, cell seeding density and geometry play an important role in experiments that require long-term spheroid monitoring. These two parameters determine the size of spheroid and total imaging time. Though the optimal spheroid size depends on the assay requirements, such as the presence or absence of hypoxic core, it usually falls within a range of 200–800 µm diameter^18,35,36^. Furthermore, the hypoxic core rapidly created by the fast-growing spheroids can interfere with phenotypic measurements. As shown previously, fast-growing spheroids produced an apoptotic signal comparable to or even greater than that induced by a drug, thus interfering with cytotoxicity measurement^22^. In such cases, the analysis of smaller spheroids may be preferable. To this end, we have demonstrated that tissueoids of various diameters could be generated with consistent size and shape controlled by adjusting the density of cell seeding. This study also showed that cells even as low as 25/well, could form tissueoids efficiently in less than 24-48 hours with high reproducibility. This feature suggests the flexibility and adaptability of the platform to generate 3D tissueoids when the sample availability is minimal.

Several previous studies have shown that 2D-cultured cells tend to overestimate the efficacies of chemotherapeutic drugs compared with 3D-cultured cells^37,38^and hence numerous anticancer drugs are eliminated during clinical development. Consistent with these studies, our present study demonstrated relative resistance to doxorubicin in 3D tissueoids as compared to 2D monolayer culture. Interestingly, differences between 3D tissueoids and 2D cultures have been observed consistently across the models. Overall, 3D tissueoids grown on AXTEX-4D™ platform 1) tended to be less sensitive to growth inhibition or apoptosis by chemotherapeutic drug (2) showed high drug resistance and viability for doxorubicin treated MCF-7 cultures than in collagen-based 3D culture. Our results are in line with the already reported literature where doxorubicin tested on different 3D models and found to be resistant^39^. However, necrotic cell death on MCF-7 tissueoids treated with high concentration of doxorubicin (10μM) on AXTEX-4D™ suggest progression of apoptotic cell into the secondary necrotic cells and needs to be evaluated further (Fig 5C). These findings speculate that a 3D-tissueoids potentially avoids the overestimation of antitumor efficacy observed in a 2D-culture and may be comparatively better than collagen-based culture. Therefore, 3D-tissueoids model could be applied as a drug testing platform to screen large compound libraries.

However, several limitations of our study warrant mention. Firstly, this study was derived from the nature of an *in vitro* model. Despite several 3D-culture systems, none of them are considered a standard method, and it is unclear which system is the most clinically relevant^7,31,40,41^. One observation is that 3D-culture of cell lines will never accurately, fully represent the tumor microenvironment *in vivo*, because the latter have interactions with stromal tissues or blood perfusions^42^. Co-culture with stromal cells or vascular cells may partially solve these issues. Secondly, patients derived tumors (PDXs) are considered a potential drug screening platform for the next generation^43,44^. Therefore, we are further investigating the application of our platform with PDXs & co-culture with stromal cells, and the process is under investigation in our laboratory.

Overall, the current study highlights the characteristics and utility of the newly developed 3D tissueoids model, which possess basic features for the 3D growth of cell lines and could be beneficial in drug screening assays. Further studies are required extensively to elucidate the mechanistic aspects of the tumor microenvironment, sensitivity towards various clinically relevant treatments with special reference to biopsy tissue from patients.

## Material and Methods

### Cell Lines

The various human cancer cell lines (MCF-7 (Breast Cancer), PC3 (Prostate cancer), A375 (Skin Melanoma), HepG2 (Liver Cancer), CHOK1 (Chinese hamster ovary), HEK-293 (Human embryonic kidney) were obtained from ATCC. All the cancer cell lines were grown in EMEM (MCF-7 and HepG2), F12K (PC3 and CHOK-1), and DMEM (A-375 and HEK-293) supplemented with 2mM glutamine (Sigma-Aldrich, St; Louis, MO, USA) and 10% FBS (Gibco) at 37°C humidified condition with 8% CO2 under static condition.

### 2D cell culture

For 2D culture, cells were seeded at around 60-80% confluency, i.e., approximately 0.8 × 106 in a 60 mm dish. At 100% confluency of cells, media was removed, and cells were washed with PBS, trypsinized, and centrifuged at 1000 rpm for 5 minutes. Finally, cells were resuspended in an appropriate volume of respective media and plated in 96 well plate (5×103), depending upon the experiment.

### 3D cell culture

Tissueoids wereformed on AXTEX-4D™ platform as described previously^17^. Briefly, cells were seeded at around 60-80% confluency, i.e., approximately 0.8 × 106 in a 60 mm dish. The cells were washed with PBS at 100% confluency, trypsinized, and centrifuged at 1000 rpm for 5 minutes. The cell suspension was made in such a way that 1 ml of respective media contain 2.5 × 10^5^ cells so that 20µl of the media contained a cell number of 5000 cells per drop. The drop was pipetted onto the inner surface of a lid filled with PBS at the bottom. After 24 −48 hours the inner lid was inverted, and the drops were poured on AXTEX-4D™ platform for tissueoids formation in the desired format and analyzed for further experiments attachment and growth. The diameter of tissueoids was measured by Biowizard software. For collagen-I 3D culture, plates were prepared by adding 50µg/ml solution of collagen prepared in 20mM acetic acid, then incubated for 1 hour at 37°C. Following that, plates were rinsed thrice with 1X PBS and were poured along with media for further experiment. 3D tissueoids were observed by using phase contrast inverted microscope (Nikon Eclipse TS100) at various magnifications.

### Scanning Electron Microscopy (SEM)

Tissueoids grown on AXTEX-4D™ platform was fixed with 2.5% glutaraldehyde and 2% paraformaldehyde in PBS, pH 7.4. After that, samples were vacuum dried for 10 mins with 0.1 mbar pressure followed by mixing of argon gas. Finally, the samples were coated with gold particles using Agar Sputter Coater (AIIMS, New Delhi, India). The coated samples were analyzed using SEM (EVO-18 Research, Zeiss) at 500, 1000, and 1500X magnification.

### Confocal Microscopy

Briefly, samples were fixed in 3.7% formaldehyde solution, permeabilized with 0.1% TRITON X-100. Phalloidin–tetramethylrhodamine B isothiocyanate conjugate solution (50µg/ml), Calcein AM (100ng/ml) and NucSpot Live 488 nuclear stain (1:1000) were used to mount cells for 5-10min at RT. For hypoxia, 2D and 3D tissueoids of MCF-7 cells were stained with HIF-1α specific antibody (Cat#ab2185). Briefly, cells were fixed with 4% PFA for 15 minutes and washed with 1X PBS. After permeabilization with 0.1% Triton-X for 4 minutes, samples were blocked with 1% BSA for 1hour at RT and next incubated with 1ug/ml primary antibody for overnight at 4°C. Finally, tissueoids were incubated with secondary antibody at 37°C for 2 hours and washing was done with PBS. Images were acquired using Nikon’s confocal microscope (A1 R HD 25) and analyzed with the NIS Elements software (Nikon Corporation, Tokyo, Japan). To avoid inter-channel mixing (405 nm, 488 nm, and 561 nm), pictures were captured separately with individual laser and were merged later.

### Treatment with clinical Drug

Both 2D and 3D cells (5000/well, n=3) were treated with doxorubicin for 48 hours in a dose dependent manner (500nM to 10000 nM). Finally, Prestoblue assay has been done as described elsewhere^45^. PrestoBlue (Thermo Fischer; A13261) is a resazurin-based solution for rapidly quantifying the metabolic active cells, providing a metric of cell viability. Briefly 1/10th volume of prestoblue reagent was directly added to the control and treated cells and incubated for 2 hours at 37°C. The metabolic rates were measured by the amount of relative fluorescence unit (RFU) at Excitation 560 and Emission 590nm using a spectrofluorometer plate reader (SPECTRA MAX GEMINI EM, Molecular Devices)

### Detection of Cell Apoptosis using Annexin V and 7-AAD

Doxorubicin treated cells were trypsinized, washed, and incubated in the dark at room temperature with Annexin V and 7-AAD for 15 mins. Following that, samples were analyzed using ACEA flow cytometry. The upper left quadrant cells (annexin V+/ 7-AAD–) as apoptotic, lower right (annexin V+/ 7-AAD+) as single positive necrotic and the upper right quadrant cells(annexin V+/ 7-AAD+) as double positive necrotic regions were defined.

## Acknowledgements

We acknowledge SAIF-Facility, AIIMS New Delhi for Scanning Microscopy facility, Confocal facility, IGIB, New Delhi for assistance and Avijit Das, Premas Biotech for his critical discussion on experimental study. We would also like to thank Guy Dauwe for his contribution and discussion on materials used in the study.

## Author Contributions

AB: Designed and planned the experiments, conducted experiments, analyzed the results, and writing. SM: Concept design, interpretation of data, and critically reviewed the manuscript. PK: Study concept, design, interpretation of data, providing scientific inputs SS: Execution and data analysis. BPD: Planning and execution of experiments. RG: Data analysis and drafted the manuscript. NM: Study concept, design, interpretation of data, providing scientific inputs and drafting of the manuscript.

## References

1 Zugazagoitia, J. et al. Current Challenges in Cancer Treatment. Clin Ther 38, 1551–1566, doi:10.1016/j.clinthera.2016.03.026 (2016).

2 Kitaeva, K. V., Rutland, C. S., Rizvanov, A. A. & Solovyeva, V. V. Cell Culture Based in vitro Test Systems for Anticancer Drug Screening. Front Bioeng Biotechnol 8, 322, doi:10.3389/fbioe.2020.00322 (2020).

3 Martin, L., Hutchens, M., Hawkins, C. & Radnov, A. How much do clinical trials cost? Nat Rev Drug Discov 16, 381–382, doi:10.1038/nrd.2017.70 (2017).

4 Hay, M., Thomas, D. W., Craighead, J. L., Economides, C. & Rosenthal, J. Clinical development success rates for investigational drugs. Nat Biotechnol 32, 40–51, doi:10.1038/nbt.2786 (2014).

5 Kapalczynska, M. et al. 2D and 3D cell cultures - a comparison of different types of cancer cell cultures. Arch Med Sci 14, 910–919, doi:10.5114/aoms.2016.63743 (2018).

6 Jensen, C. & Teng, Y. Is It Time to Start Transitioning From 2D to 3D Cell Culture? Front Mol Biosci 7, 33, doi:10.3389/fmolb.2020.00033 (2020).

7 Breslin, S. & O’Driscoll, L. Three-dimensional cell culture: the missing link in drug discovery. Drug Discov Today 18, 240–249, doi:10.1016/j.drudis.2012.10.003 (2013).

8 Nyga, A., Cheema, U. & Loizidou, M. 3D tumour models: novel in vitro approaches to cancer studies. J Cell Commun Signal 5, 239–248, doi:10.1007/s12079-011-0132-4 (2011).

9 Sutherland, R. M. Cell and environment interactions in tumor microregions: the multicell spheroid model. Science 240, 177–184, doi:10.1126/science.2451290 (1988).

10 Benien, P. & Swami, A. 3D tumor models: history, advances and future perspectives. Future Oncol 10, 1311–1327, doi:10.2217/fon.13.274 (2014).

11 Kim, J. E., Kim, S. H. & Jung, Y. Current status of three-dimensional printing inks for soft tissue regeneration. Tissue Eng Regen Med 13, 636–646, doi:10.1007/s13770-016-0125-8 (2016).

12 K. Luo, Y. Y.Z. Shao. Physically crosslinked biocompatible silk-fibroin-based hydrogels with high mechanical performance. Adv. Funct. Mater., 26, 872–880 (2016).

13 Jongpaiboonkit, L. et al. An adaptable hydrogel array format for 3-dimensional cell culture and analysis. Biomaterials 29, 3346–3356, doi:10.1016/j.biomaterials.2008.04.040 (2008).

14 Cai, S., Xu, H., Jiang, Q. & Yang, Y. Novel 3D electrospun scaffolds with fibers oriented randomly and evenly in three dimensions to closely mimic the unique architectures of extracellular matrices in soft tissues: fabrication and mechanism study. Langmuir 29, 2311–2318, doi:10.1021/la304414j (2013).

15 Knight, E. & Przyborski, S. Advances in 3D cell culture technologies enabling tissue -like structures to be created in vitro. J Anat 227, 746–756, doi:10.1111/joa.12257 (2015).

16 Kundu P, M. N., Das A, Mazumdar S, Baru A. Device and Methods for Multidimentional Cell Culture. (US 20200326330 A1 2020).

17 Ambica Baru, S. S., Biswa Pratim Das Purkayastha, Sameena Khan, Saumyabrata Mazumder, Reeshu Gupta, Prabuddha Kumar Kundu, Nupur Mehrotra Arora. AXTEX-4D™: A Novel 3D ex vivo platform for preclinical investigations of immunotherapy agents. BioRxiv, doi:10.1101/2020.11.05.369751 (2020).

18 Vinci, M. et al. Advances in establishment and analysis of three-dimensional tumor spheroid-based functional assays for target validation and drug evaluation. BMC Biol 10, 29, doi:10.1186/1741-7007-10-29 (2012).

19 Vinci, M., Box, C. & Eccles, S. A. Three-dimensional (3D) tumor spheroid invasion assay. J Vis Exp, e52686, doi:10.3791/52686 (2015).

20 Silva, I. A. et al. Aldehyde dehydrogenase in combination with CD133 defines angiogenic ovarian cancer stem cells that portend poor patient survival. Cancer Res 71, 3991–4001, doi:10.1158/0008-5472.CAN-10-3175 (2011).

21 Raghavan, S. et al. Formation of stable small cell number three-dimensional ovarian cancer spheroids using hanging drop arrays for preclinical drug sensitivity assays. Gynecol Oncol 138, 181–189, doi:10.1016/j.ygyno.2015.04.014 (2015).

22 Mittler, F. et al. High-Content Monitoring of Drug Effects in a 3D Spheroid Model. Front Oncol 7, 293, doi:10.3389/fonc.2017.00293 (2017).

23 Shi, W. et al. Facile Tumor Spheroids Formation in Large Quantity with Controllable Size and High Uniformity. Sci Rep 8, 6837, doi:10.1038/s41598-018-25203-3 (2018).

24 Chan, H. F. et al. Rapid formation of multicellular spheroids in double-emulsion droplets with controllable microenvironment. Sci Rep 3, 3462, doi:10.1038/srep03462 (2013).

25 McMillan, K. S., McCluskey, A. G., Sorensen, A., Boyd, M. & Zagnoni, M. Emulsion technologies for multicellular tumour spheroid radiation assays. Analyst 141, 100–110, doi:10.1039/c5an01382h (2016).

26 Sabhachandani, P. et al. Generation and functional assessment of 3D multicellular spheroids in droplet based microfluidics platform. Lab Chip 16, 497–505, doi:10.1039/c5lc01139f (2016).

27 Chen, Y. et al. Metabolic profiling of normal hepatocyte and hepatocellular carcinoma cells via H nuclear magnetic resonance spectroscopy. Cell Biol Int 42, 425–434, doi:10.1002/cbin.10911 (2018).

28 Abu El Maaty, M. A., Alborzinia, H., Khan, S. J., Buttner, M. & Wolfl, S. 1,25(OH)2D3 disrupts glucose metabolism in prostate cancer cells leading to a truncation of the TCA cycle and inhibition of TXNIP expression. Biochim Biophys Acta Mol Cell Res 1864, 1618–1630, doi:10.1016/j.bbamcr.2017.06.019 (2017).

29 Walker, M., Rizzuto, P., Godin, M. & Pelling, A. E. Structural and mechanical remodeling of the cytoskeleton maintains tensional homeostasis in 3D microtissues under acute dynamic stretch. Sci Rep 10, 7696, doi:10.1038/s41598-020-64725-7 (2020).

30 Liverani, C. et al. A biomimetic 3D model of hypoxia-driven cancer progression. Sci Rep 9, 12263, doi:10.1038/s41598-019-48701-4 (2019).

31 Weigelt, B., Ghajar, C. M. & Bissell, M. J. The need for complex 3D culture models to unravel novel pathways and identify accurate biomarkers in breast cancer. Adv Drug Deliv Rev 69-70, 42–51, doi:10.1016/j.addr.2014.01.001 (2014).

32 Hirt, C. et al. Bioreactor-engineered cancer tissue-like structures mimic phenotypes, gene expression profiles and drug resistance patterns observed “in vivo”. Biomaterials 62, 138–146, doi:10.1016/j.biomaterials.2015.05.037 (2015).

33 Unger, C. et al. Modeling human carcinomas: physiologically relevant 3D models to improve anti-cancer drug development. Adv Drug Deliv Rev 79-80, 50–67, doi:10.1016/j.addr.2014.10.015 (2014).

34 Tibbitt, M. W. A.K. S.. Hydrogels as extracellular matrix mimics for 3D cell culture. Biotechnol Bioeng 103, 655–663 (2009).

35 Fischbach, C. et al. Engineering tumors with 3D scaffolds. Nat Methods 4, 855–860, doi:10.1038/nmeth1085 (2007).

36 Ivanov, D. P. et al. Multiplexing spheroid volume, resazurin and acid phosphatase viability assays for high-throughput screening of tumour spheroids and stem cell neurospheres. PLoS One 9, e103817, doi:10.1371/journal.pone.0103817 (2014).

37 Herter, S. et al. A novel three-dimensional heterotypic spheroid model for the assessment of the activity of cancer immunotherapy agents. Cancer Immunol Immunother 66, 129–140, doi:10.1007/s00262-016-1927-1 (2017).

38 Bleul, C. C., Fuhlbrigge, R. C., Casasnovas, J. M., Aiuti, A. & Springer, T. A. A highly efficacious lymphocyte chemoattractant, stromal cell-derived factor 1 (SDF-1). J Exp Med 184, 1101–1109, doi:10.1084/jem.184.3.1101 (1996).

39 Lovitt, C. J., Shelper, T. B. & Avery, V. M. Doxorubicin resistance in breast cancer cells is mediated by extracellular matrix proteins. BMC Cancer 18, 41, doi:10.1186/s12885-017-3953-6 (2018).

40 Rimann, M. & Graf-Hausner, U. Synthetic 3D multicellular systems for drug development. Curr Opin Biotechnol 23, 803–809, doi:10.1016/j.copbio.2012.01.011 (2012).

41 Lovitt, C. J., Shelper, T. B. & Avery, V. M. Advanced cell culture techniques for cancer drug discovery. Biology (Basel) 3, 345–367, doi:10.3390/biology3020345 (2014).

42 Imamura, Y. et al. Comparison of 2D- and 3D-culture models as drug-testing platforms in breast cancer. Oncol Rep 33, 1837–1843, doi:10.3892/or.2015.3767 (2015).

43 Ghosh, G., Lian, X., Kron, S. J. & Palecek, S. P. Properties of resistant cells generated from lung cancer cell lines treated with EGFR inhibitors. BMC Cancer 12, 95, doi:10.1186/1471-2407-12-95 (2012).

44 Kurokawa, M., Ise, N., Omi, K., Goishi, K. & Higashiyama, S. Cisplatin influences acquisition of resistance to molecular-targeted agents through epithelial-mesenchymal transition-like changes. Cancer Sci 104, 904–911, doi:10.1111/cas.12171 (2013).

45 Tian, T. et al. Rac1 is a novel therapeutic target in mantle cell lymphoma. Blood Cancer J 8, 17, doi:10.1038/s41408-018-0052-0 (2018).

